# NECAP antagonizes light-induced Rhodopsin-1 internalization to promote photoreceptor homeostasis

**DOI:** 10.64898/2026.06.01.729193

**Authors:** Huai-Wei Huang, Michal Tyrlik, Po-Wen Huang, Adam I. Yeo, Ah-Ram Kim, Norbert Perrimon, Chen-Yuan Tseng, Hugo J. Bellen, Hyung Don Ryoo

## Abstract

AP-2 is a key mediator of clathrin-mediated endocytosis (CME) that internalizes cargo from the plasma membrane. NECAP proteins physically bind phosphorylated AP-2 during CME, but their roles during endocytosis remain unresolved, with conflicting reports about their function. Here, we report that *Drosophila NECAP* is dispensable for development, but antagonizes light-dependent Rhodopsin-1 (Rh1) internalization, a process that occurs through AP-2-mediated endocytosis. Specifically, loss of *Drosophila NECAP* causes excessive light-dependent Rh1 internalization and an age-related retinal degeneration that can be rescued by photoreceptor-specific expression of a wild-type transgenic *NECAP.* A *Drosophila NECAP* mutant transgene, equivalent to a canine *NECAP1* variant associated with retinal atrophy, failed to rescue the *NECAP* loss-of-function phenotype in the eye. Furthermore, overexpression of wild-type *NECAP* suppressed massive Rh1 internalization in a distinct *Drosophila* model of light-dependent Rh1 endocytosis and retinal degeneration. These results establish NECAP as a negative regulator of light-dependent Rh1 internalization essential for photoreceptor survival.

## Introduction

Clathrin-mediated endocytosis (CME) is a major membrane trafficking pathway that internalizes proteins and other macromolecules from the plasma membrane, regulating diverse biological processes including nutrient uptake, receptor signaling, synaptic vesicle recycling, and cell polarity (Mettlen et al. 2018; Kirchhausen, Owen, and Harrison 2014). Central to CME is the heterotetrameric adaptor protein complex 2 (AP-2), composed of α, β2, μ2, σ2 subunits. AP-2 serves as a critical coordinator of CME while undergoing a series of changes in localization, conformation, and interactions with accessory factors. Therefore, a better understanding of the AP-2’s interaction partners could provide important biological and pathological insights associated with CME.

During the initiation stages of CME, AP-2 is recruited to the plasma membrane in a closed conformation through its interaction with PhosphatidylInositol-4,5-bisPhosphate (PIP_2_) (Collins et al. 2002; Höning et al. 2005). Subsequently, membrane-associated muniscin proteins trigger AP-2 to assume an active open conformation (Henne et al. 2010; Hollopeter et al. 2014; Umasankar et al. 2014), leading to a stable membrane interaction, recruitment of clathrin, and membrane-embedded cargo (Owen and Evans 1998; Kadlecova et al. 2017; Jackson et al. 2010; Kelly et al. 2014). AP-2 undergoes multiple phosphorylation events, and perhaps the best-characterized is the one at Thr156 of AP-2mu (in human and mice) that occurs when the complex adopts an open conformation after associating with the membrane and cargo (Höning et al. 2005; Olusanya et al. 2001; Pauloin and Thurieau 1993).

NECAP (adaptin-ear-binding coat-associated protein) 1 and 2 in mammals and *C. elegans* ncap-1 are proteins that bind to AP-2 phosphorylated at Thr156 (Ritter et al. 2013; Beacham 2018; Partlow 2019; Wrobel et al. 2019). Structural studies have shown that the N-terminal 133-residue PHear domain directly binds to AP-2mu phosphorylated at Thr156 (Partlow, Baker et al. 2019, Wrobel, Kadlecova et al. 2019). In addition, mammalian NECAP proteins contain a C-terminal WVQF motif that binds to AP-2’s α subunit (Ritter et al. 2003; Praefcke et al. 2004). This WVQF motif is absent in the *C. elegans* ncap-1, but the *Drosophila* homolog has high sequence conservation with mammalian NECAPs throughout the coding sequence, including the PHear and the WVQF motifs (Ritter et al. 2003). The role of NECAP during CME remains unresolved with conflicting reports about its function. For example, some studies have concluded that mammalian NECAP proteins stimulate CME after binding to phospho-AP-2 (Ritter et al. 2013; Wrobel et al. 2019), whereas others report an antagonistic role during CME (Beacham 2018; Partlow 2019). Additionally, NECAP2 has been shown to function independently of clathrin-coated vesicles (CCVs) by regulating the AP-1 adaptor complex to promote recycling of client proteins from the endosome to the plasma membrane (Chamberland et al. 2016). Furthermore, *NECAP* loss-of-function phenotypes in animal models remain unknown, aside from the reported genetic interaction between *C. elegans ncap-1* and *fcho-1* (encoding a muniscin) (Beacham 2018).

Among the various CME cargo proteins are *Drosophila* Rhodopsin-1 (Rh1), encoded by the *ninaE* gene, a light sensing protein of the visual signaling cascade. (Montell 2012; O’Tousa 1985; Zuker 1985). Rh1 is expressed at high levels in R1 to R6 photoreceptors in the *Drosophila* retina, primarily in the rhabdomere, a specialized apical membrane compartment composed of tightly packed microvilli. Upon light activation, Rh1 at the rhabdomere binds Arrestin and undergoes AP-2-mediated endocytosis (Satoh and Ready 2005; Orem, Xia, and Dolph 2006). Rh1-containing CCVs traffic to endosomes, where Rh1 is either transported to the lysosome for degradation (Orem, Xia, and Dolph 2006; Satoh and Ready 2005; Mu et al. 2019) or recycled back to the rhabdomeres (Wang et al. 2014). Genetic conditions that cause excessive internalization of light-activated Rh1 can cause retinal degeneration (Kiselev et al. 2000; Orem, Xia, and Dolph 2006; Chincore 2009; Xu and Wang 2016; Dourlen et al. 2012; Acharya et al. 2008; Huang 2021; Wang et al. 2014). This retinal degeneration can be suppressed by blocking Rh1 endocytosis (Wang et al. 2014; Dourlen et al. 2012), or by enhancing Rh1 recycling from endosomes to the apical membranes (Wang et al. 2014), suggesting that high levels of Rh1 in endosomes contribute to photoreceptor degeneration.

In mammalian rod cells, phototransduction occurs in the outer segment, containing stacks of membrane discs packed with Rhodopsin. There is no evidence that light-activated Rhodopsins undergo endocytosis in the outer segment, but mammalian Rhodopsin variants with high affinity for Arrestin undergo internalization in the inner segment of rod cells, contributing to severe forms of Retinitis Pigmentosa (RP) (Chuang et al. 2004; Chen et al. 2006). Whether mammalian NECAP has any roles in Rhodopsin homeostasis remains unknown. Interestingly, a *NECAP1* variant has been reported in a canine breed predisposed to retinal atrophy (Hitti et al. 2019), but the cellular/molecular role of *NECAP1* in the eye has not been examined.

Here, we report the role of *CG9132*, a single *Drosophila* homolog of *NECAP1* and *2*, in photoreceptor homeostasis (henceforth referred to as *NECAP*). We show that *Drosophila NECAP* loss-of-function mutants survive to adulthood, suggesting that *NECAP* is not essential for all clathrin-mediated endocytosis in this organism. We further find massive internalization of light-dependent Rh1 and retinal degeneration in *NECAP* mutants, which can be rescued by a photoreceptor-specific expression of transgenic human *NECAP1, NECAP2,* or *Drosophila NECAP.* Furthermore, *Drosophila NECAP* variants equivalent to those associated with human and canine diseases behave as loss-of-function alleles with dominant negative properties. Overexpression of wild type *NECAP* is sufficient to suppress Rh1 internalization in other models of clathrin-mediated Rh1 endocytosis. Taken together, these findings establish NECAP as a negative regulator of Rh1 internalization essential for photoreceptor survival.

## Results

### *Drosophila NECAP* loss causes light-dependent retinal degeneration

The *Drosophila* genome carries a single uncharacterized homolog of mammalian *NECAP1* and *NECAP2* on the X chromosome, *CG9132*. It encodes two protein isoforms generated by alternative splicing. The longer isoform is of 246 amino acid residues, and the shorter isoform is of 204 amino acids (Figure S1a). To understand the biological role of *Drosophila NECAP*, we examined two loss-of-function alleles: One is a knockout line we generated through CRISPR-Cas9-mediated deletion (*NECAP^KO^*), replacing most of the NECAP coding sequence with a transgene encoding a fluorescent marker (3xP3-RFP) (Figure 1a; Figure S1b). Specifically, the shorter protein-coding sequence is completely lost, while only the first 27 N-terminal amino acid residues of the longer isoform remain, deleting all known functional domains including the PHear domain (a.a. 1 – 130) and the C-terminal WVQF motif.

**Figure 1:**
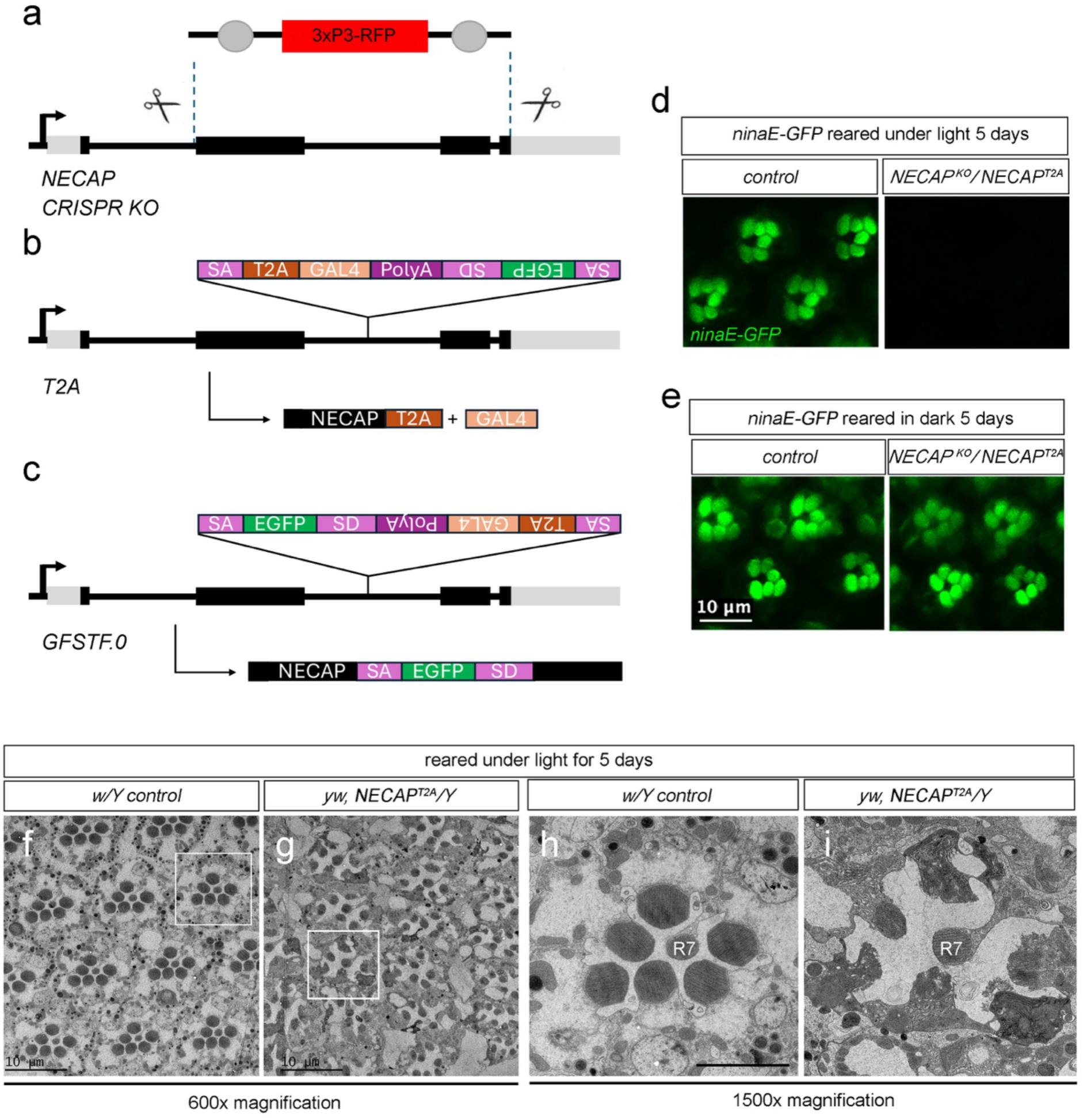
The *NECAP* mutants show abnormal *ninaE* expression and photoreceptor morphology. Because eye pigments could affect the light-dependent phenotypes, all data were generated in white eyed flies. Specifically, the *NECAP^CRISPRKO^* line was in a *w^1118^*background, while the *NECAP^T2A^* was in the *yw* genetic background. (a-c) Schematic diagrams of the *NECAP* alleles. The coding sequences are in black rectangles, and the UTRs are in gray. (a) The knockout (KO) allele. The dotted lines indicate sites targeted by the gRNAs for deletion. The deleted NECAP coding sequence was replaced with a 3xP3-RFP transgene. (b) The *T2A* allele. (c) The *GFSTF.0* allele. The expected protein products are depicted below the genomic locus. SA indicates a splice acceptor, and SD indicates a splice donor. The T2A element truncates the encoded protein, releasing the Gal4 protein in the *T2A* allele. (d, e) Photoreceptor integrity and Rh1 expression as assessed through the *ninaE-GFP* reporter (*ninaE* encodes the Rh1 protein). In healthy control retina (genotype: *yw, ninaE-GFP/w*), GFP-expressing photoreceptors are clustered in a trapezoidal pattern in each ommatidium. Note that GFP fluorescence is absent in the *yw, ninaE-GFP, NECAP^T2A^/w, NECAP^KO^* flies reared under light (d). Dark reared *yw, ninaE-GFP, NECAP^T2A^/w, NECAP^KO^* retina have GFP-expressing photoreceptors similar to control flies (e). (f- i) Transmission electron microscopy images of eyes from five-day-old adult flies reared under constant light. Control *w^1118^/Y* (f, h) was compared with *yw, NECAP^T2A^/Y* (g, i). 600x magnification images show arrays of ommatidia (f, g), while the 1500x magnification images show a single ommatidium with seven rhabdomeres (h, i). The magnified ommatidia are from the inset shown in (f, g). R7 rhabdomere, which does not express Rh1, remains intact in *yw, NECAP^T2A^/Y*. But the other rhabdomeres (R1 to R6), which all express Rh1, are severely disrupted in the *yw, NECAP^T2A^/Y* retina (i). The scale bar in e represents panels d, e and indicates 10 μm. The scale bar in h, represents both h and i to indicate 5 μm.

Two other mutants were created by conversion of a *Minos*-based transposon, MiMIC (MI 15073), inserted in NECAP using the double header strategy (Li-Kroeger et al. 2018). This allows the MiMIC element to be converted to a *GFP*-tagged protein trap and/or a *T2A-GAL4* gene trap (Nagarkar-Jaiswal et al., 2015)(Figure 1b, c). The *T2A-GAL4* gene trap contains a splice acceptor sequence followed by *T2A-Gal4-PolyA* inserted in the second intron of this gene (Figure 1b; Figure S1c). The T2A-Gal4-polyA sequence is inserted after NECAP’s 160^th^ amino acid residue. The insertion will generate a *NECAP-T2A-Gal4* transcript, which will result in a truncated NECAP protein and an active Gal4 present in *NECAP* expressing cells upon translation. We confirmed the deletion of the targeted region in the *NECAP^KO^* with genomic PCR (Figure S1c) and performed RT-PCR to confirm the absence of *NECAP* transcripts downstream of the insertion site in *NECAP^T2A^/Y* males (Figure S1e). We assessed the expression pattern of *NECAP* by taking advantage of the Gal4 produced by *NECAP^T2A^* that can drive *UAS-tdTomato* expression. The tdTomato fluorescence was detected throughout the adult body, including the adult photoreceptors (Figure S2). All the mutant alleles produce viable adults without obvious morphological defects at the time of adult eclosion, which contrasts with the early developmental lethality associated with the loss of AP-2 subunits (González-Gaitán and Jäckle 1997; Windler and Bilder 2010).

As mammalian *NECAP1* and *NECAP2* are implicated in endocytosis regulation, and since light-activated Rh1 undergoes degradation through endocytosis, we examined *ninaE-GFP*, a transgene with a GFP reporter fused to the Rh1’s coding sequence to track Rh1 levels in live fly photoreceptors (Huang, Xie, and Wang 2015). In control *ninaE-GFP* flies, the GFP fluorescence is visible in all six Rh1-expressing photoreceptors (specifically, R1 to R6 cells) that are arranged in a trapezoidal pattern in each ommatidium. In *ninaE-GFP, NECAP^T2A^/NECAP^KO^* flies reared under constant light for five days, the GFP fluorescence was lost (Figure 1d). On the other hand, equivalent flies reared in the dark had *ninaE-GFP* expression comparable to controls (Figure 1e), indicating that the retina phenotype of *NECAP^T2A^* is light-dependent.

The loss of *ninaE-GFP* signal could be due to either reduced *ninaE-GFP* expression or retinal degeneration. To examine the integrity of the retina, we subjected five-day-old flies reared under constant light through transmission electron microscopy. Control flies (*w^1118^* males) maintained a regular array of ommatidia, each with seven rhabdomeres arranged in a trapezoidal pattern (Figure 1f). By contrast, *yw, NECAP^T2A^/Y* males reared under equivalent conditions had disorganized ommatidia. Many had fewer than seven rhabdomeres in an ommatidium, and the ommatidia were no longer arranged in a regular trapezoidal pattern. In addition, there were numerous vacuoles in between the ommatidia of *yw, NECAP^T2A^/Y* males (Figure 1g). Ommatidia with seven rhabdomeres organized in a trapezoidal pattern are evident in higher magnification images of wild type flies. R7 photoreceptor at the center, which does not express Rh1, is surrounded by R1 to R6 photoreceptors that express Rh1 (Figure 1h). In the *yw, NECAP^T2A^/Y* retina, some ommatidia had intact R7 cells surrounded by degenerating R1 to R6 photoreceptors, as indicated by the more electron-dense (darker) cytoplasms and the lack of normal rhabdomeres (Figure 1i). To further corroborate the results, we examined the retina of *yw, NECAP^T2A^/w, NECAP^KO^* adult flies reared under constant light for 5 days. Visualization of the photoreceptor rhabdomeres with phalloidin labeling revealed a well-organized array of ommatidia in controls but not in the *NECAP* homozygous retina (Figure S3). These results indicate that *NECAP* is essential to maintain the integrity of photoreceptors that express Rh1 when reared under light.

### Structural conservation of *Drosophila* NECAP as modeled by AlphaFold

While the crystallography and cryoEM-based studies could not visualize mammalian NECAP’s C-terminal WVQF interacting with AP-2, the reported structures of NECAP1 and 2 show the PHear domain directly interacting with the AP-2mu subunit phosphorylated at Thr156. (Wrobel, Kadlecova et al. 2019; Partlow, Baker et al. 2019; Figure 2a). We modeled the interaction between NECAP and AP-2mu using AlphaFold 3 to assess *Drosophila* NECAP’s structural conservation (Abramson et al. 2024). We specifically applied integrated Local Interaction Score (iLIS), a metric recently used to survey the *Drosophila* protein-protein interactions at a proteomic scale (Kim 2026). iLIS combines confidence scores from both the broader interaction interface between two proteins and the core physical contacts, and an iLIS value above 0.223 supports interactions with high confidence. When the modeling was performed with the unphosphorylated form of AP-2, the NECAP PHear domain/AP-2mu interaction was not supported, with an iLIS value of 0 (Figure 1b, e). By contrast, modeling with the Thr156-phosphorylated form of mammalian AP-2 predicted a high-confidence interaction with the PHear domain of NECAP-2 (Figure 2c, d), consistent with the published structural studies. *Drosophila* AP-2mu phosphorylated at the equivalent Thr154 residue was also predicted to interact with the *Drosophila* NECAP’s PHear domain (Figure 2f, g), supporting a phylogenetically conserved structure of NECAP.

**Figure 2:**
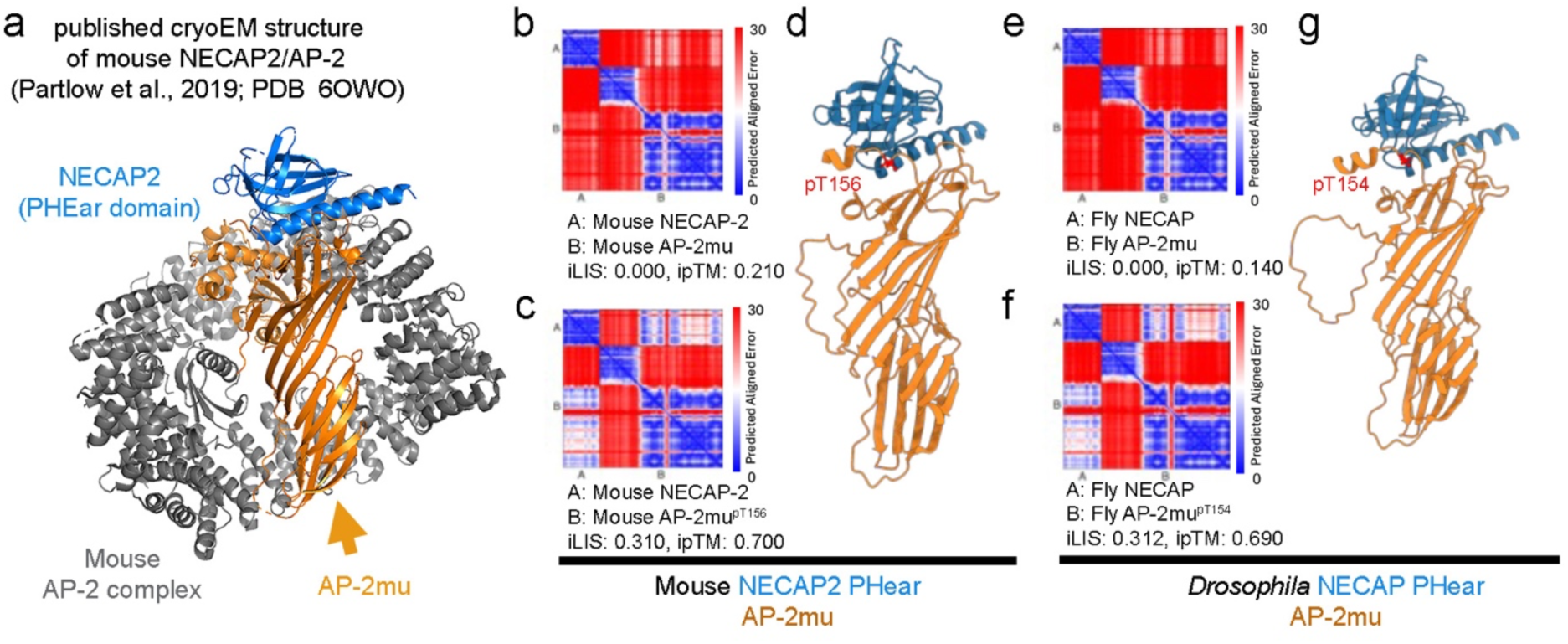
Phylogenetic conservation of the predicted NECAP PHear domain structure. In all structures, AP-2mu is shown in orange, and NECAP PHear domains are in marine blue. (a) A previously published cryoEM structure of the mouse NECAP2/AP-2 complex (PDB 6OWO). The subunits of AP-2 other than AP-2mu are shown in gray. (b – g) AlphaFold 3 predictions of the NECAP PHear domain/AP-2mu interaction. (b) Predicted aligned error (PAE) map for mouse NECAP2 and unphosphorylated AP-2mu. The iLIS value is 0, not supportive of a physical interaction between the two proteins. (c, d) Predicted interaction between mouse NECAP2 and AP-2mu phosphorylated at Thr156: PAE map (iLIS = 0.310 that supports the interaction) (c) and the predicted structure (d) consistent with the published cryoEM structure (a). (e) PAE map for *Drosophila* NECAP and the unphosphorylated AP-2, with iLIS value of 0 that does not support physical interaction. (f, g) *Drosophila* NECAP and AP-2mu phosphorylated at Thr154, supporting a predicted interaction. PAE map (iLIS = 0.312) (f) and the predicted structure (g). Phosphorylated Thr residues (mouse pT156 and *Drosophila* pT154) are colored in red.

### *NECAP* loss reduces Rh1 levels, which can be rescued by expressing wild type *Drosophila* and human *NECAP* transgenes

Because light-dependent degradation of Rh1 occurs through AP-2-mediated endocytosis (Alloway, Howard, and Dolph 2000; Kiselev et al. 2000; Orem, Xia, and Dolph 2006; Satoh and Ready 2005), we examined if *NECAP* loss affects Rh1 levels. Compared to control (*w^1118^/Y*) fly extracts, *yw, NECAP^T2A^/Y* males reared under constant light for five days had significantly reduced Rh1 as assessed through western blot. Expressing *NECAP* by introducing *UAS-NECAP* into the *yw, NECAP^T2A^/Y* male background restored total Rh1 levels closer to control flies (Figure 3b lanes 1-3, 3c). Rh1 levels were also rescued when human *NECAP1* (*hNECAP1*) or *NECAP2* (*hNECAP2*) were expressed by introducing *UAS-hNECAP1* or *-hNECAP2* into the *yw, NECAP^T2A^/Y* background (Figure 3b lanes 6 and 7, 3c). When these flies were reared in the dark, Rh1 levels did not decrease in *yw, NECAP^T2A^/Y* males, indicating that *NECAP* specifically regulates Rh1 exposed to light (Figure 3d, e).

**Figure 3:**
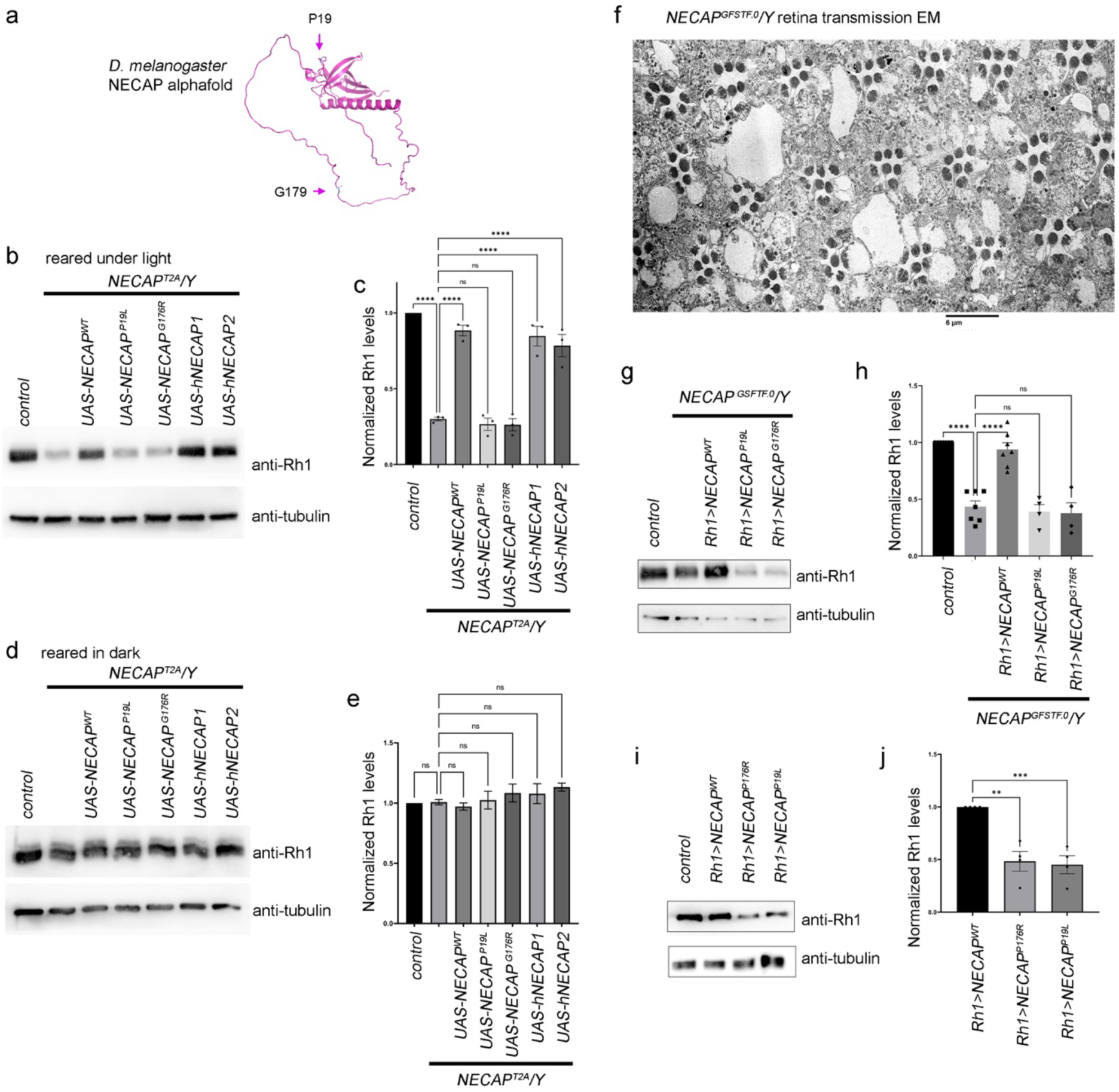
*NECAP* mutants have reduced Rh1 levels, which are rescued by expressing wild type *NECAP* transgenes but not by variants associated with diseases. For all analyses of *NECAP* mutants, hemizygous males in the *yw* genetic background were examined. (a) The position of the amino acids substituted in the disease-associated variants. (b) Western blot of Rh1 (top gel) and the loading control (bottom, anti-tubulin) from five-day-old adult male flies reared under light. (lane 1) control *w^1118^* samples. (lanes 2-7) *NECAP^T2A^/Y* samples, either without expressing any transgene (lane 2), or with the indicated *UAS-* rescue transgenes (lanes 3-7). (c) Quantification of the relative Rh1 band intensities (Rh1 band intensities divided by anti-tubulin band intensities) of the indicated genotypes. Note the reduction of Rh1 in *NECAP^T2A^/Y* flies, which is rescued specifically by expressing wild type *Drosophila NECAP,* and human *NECAP1 and NECAP2. NECAP^P19L^* and *NECAP^G176R^*are mutant transgenes equivalent to those associated with mammalian diseases. (d, e) Equivalent analysis to (c, e), but with flies reared in the dark. Note that the reduction of Rh1 in *NECAP^T2A^/Y* only occurs in flies reared under light. (f) Transmission EM image of the retina of a five day-old *yw, NECAP^GFSTF.0^/Y* hemizygous male reared under light. Note the presence of seven rhabdomeres in most ommatidia. (g, h) Western blot of Rh1 (top gel) and the loading control (bottom, anti-tubulin) from five-day-old adult male flies reared under light (g). (h) Quantification of the relative Rh1 levels (normalized to tubulin bands). Rh1 is reduced in *NECAP^GFSTF.0^/Y* male samples, which is rescued by wild type *NECAP* transgenes, but not by those equivalent to the mammalian disease-associated variants. (i, j) *NECAP^P19L^*and *NECAP^G176R^* transgenes have dominant negative properties. Shown are western blots (i) and the quantification of the relative Rh1 band intensities after normalization to tubulin (j). Overexpression of the *NECAP^P19L^*and *NECAP^G176R^* transgenes using the *Rh1-Gal4* driver, in a *NECAP wild type* background, reduces Rh1 levels. ANOVA followed by a multiple comparisons test was used for all statistical analyses. N.s. = not significant. ** p < 0.005, *** p < 0.0005, **** p < 0.00005. The scale bar in (f) represents 6 μm.

Since *yw, NECAP^T2A^/Y* males show light-dependent retinal degeneration, it remained unclear whether Rh1 is reduced in the mutant due to photoreceptor degeneration or to Rh1 protein degradation. To uncouple protein degradation from photoreceptor cell death, we employed the second line, *NECAP^GFSTF.0^*. This line has a GFP-carrying artificial exon after the 160^th^ amino acid residue of the NECAP protein (Figure 1c). Unlike the *T2A* allele, this construct has a splice acceptor and a splice donor which results in the insertion of a functional exon which encodes GFP in-frame with the full-length protein. We find that this tag partially disrupts the protein function and is a hypomorphic allele. The hemizygous males reared under light for 5 days had retinas with vacuoles, but most photoreceptors maintained intact rhabdomeres, with seven rhabdomeres in each ommatidium visible in a trapezoidal pattern in transmission EM images (Figure 3f). Although the mutant flies had intact rhabdomeres, Rh1 levels were significantly reduced when assessed through western blots, and expression of a wild type Rh1 transgene with the *Rh1-Gal4* restored Rh1 levels in the *yw, NECAP^GFSTF.0^/Y* males (Figure 3g, h).

### *NECAP* variants associated with disease fail to rescue Rh1 reduction in the fly *NECAP* mutants

We assessed whether two different variant *NECAP* transgenes can rescue the *NECAP* mutant phenotype. One variant was introduced in a *UAS-* transgene that drives the *Drosophila NECAP* allele*, P19L*, which corresponds to the human *NECAP1* P23L variant (68C>T) detected in a heterozygous carrier with developmental and epileptic encephalopathy. This variant is listed in ClinVar as a variant of uncertain significance (VUS) and affects a conserved amino acid residue in the PHear domain of NECAP (Figure S1a; Figure 3a). The pathogenicity of this variant is unclear because other *NECAP1* variants associated with early infantile epileptic encephalopathy are found in homozygous states (Alazami et al. 2014; Mizuguchi et al. 2019; Chouery et al. 2022). In addition to *P19L*, we generated a *UAS-*transgene driving the *Drosophila NECAP G176R* variant (Figure S1a; Figure 3a), which is equivalent to the *G182R* variant of canine *NECAP1* associated with retinal atrophy in certain breeds (Hitti et al. 2019). Unlike the wild type *UAS-NECAP* transgenes, both variant *NECAP* transgenes failed to rescue Rh1 levels in *yw, NECAP^T2A^/Y* males reared under light (Figure 3b, c). We also examined their ability to rescue Rh1 reduction in the weaker allele*, NECAP^GFSTF.0^/Y* flies, by driving the transgenes with the photoreceptor-specific *Rh1-Gal4* driver. The two mutant variants also failed to rescue the *NECAP^GFSTF.0^* phenotype (Figure 3g, h) whereas the wt allele fully rescues the phenotype.

### The *NECAP* alleles *P19L* and *G176R* have dominant negative properties

Since human *NECAP1 P23L* variant was detected in a heterozygous individual, we assessed if the equivalent *Drosophila* allele has dominant negative properties. We specifically overexpressed *NECAP^P19L^* in an otherwise *NECAP wild type* background using the *Rh1-Gal4* driver. Total Rh1 levels were lower than in controls. Similarly, overexpressing *NECAP^G176R^* also reduced total Rh1 as assessed through western blot (Figure 3i, j), supporting that they are dominant negative variants.

### Cytoplasmic Rh1 puncta partially co-localize with an endosome marker and accumulate in the photoreceptors of *NECAP* hypomorphic flies reared under light, which is suppressed by wild type NECAP transgene expression

In wild type flies, most Rh1 are detected within the rhabdomeres, but conditions that cause excessive Rh1 internalization typically show Rh1 in the endosomes that appear as cytoplasmic puncta (Figure 4a, b). Using immunohistochemistry, we examined Rh1’s subcellular localization in three-day-old *yw, NECAP^GFSTF.0^/Y* males, a time point that precedes retinal degeneration (Figure 3f). In wild type flies, most Rh1 co-localized with fluorescent phalloidin, which labels the rhabdomere. By contrast, *yw, NECAP^GFSTF.0^/Y* flies had considerable anti-Rh1 signals outside of the rhabdomeres, consistent with excessive Rh1 internalization into the cell body (Figure 4c, central panels). This phenotype was light-dependent, as flies reared in the dark showed Rh1 mostly localized within the rhabdomeres (Figure 4c, right).

**Figure 4:**
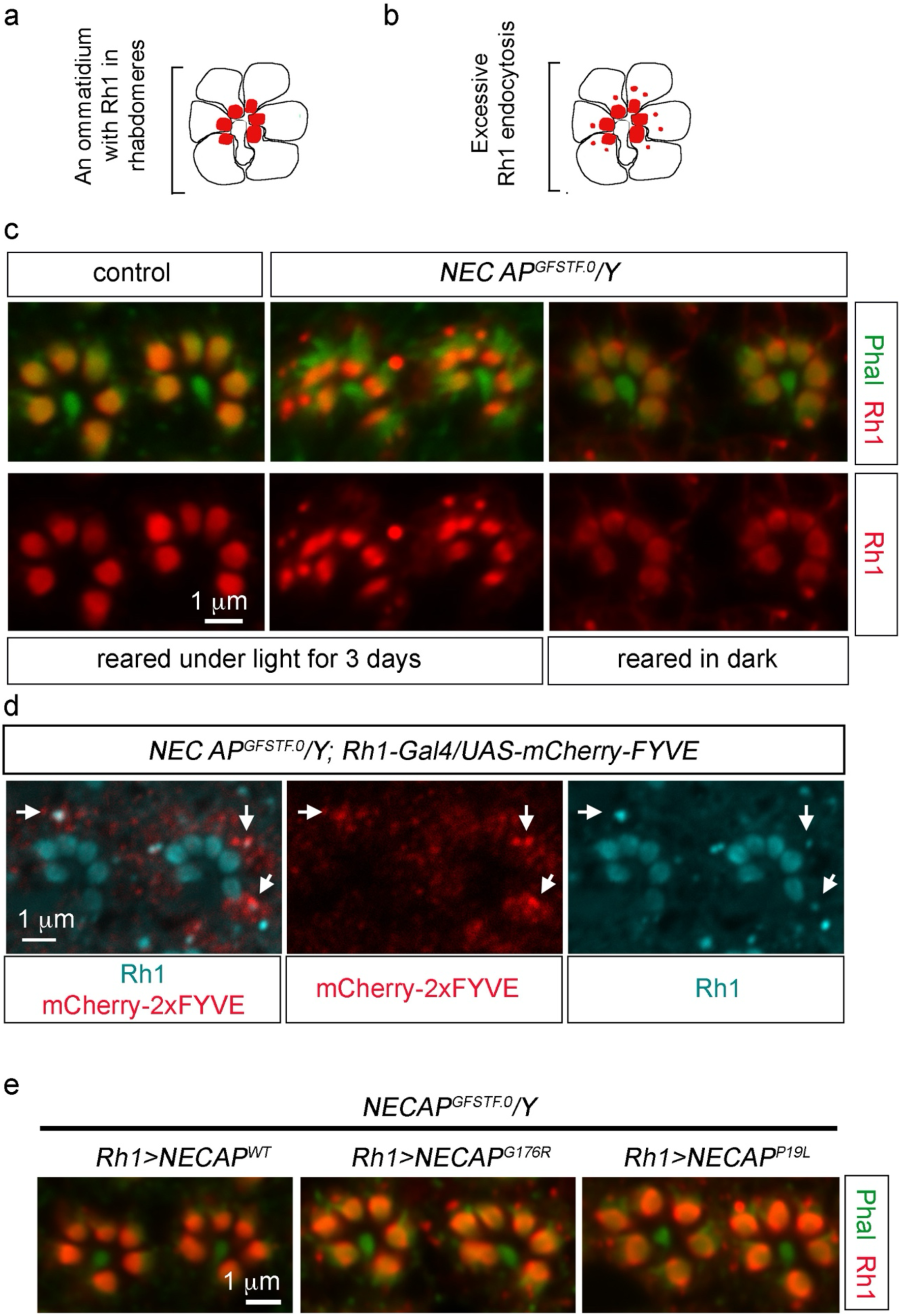
Endosomal Rh1 puncta appear in *NECAP^GFSTF.0^/Y* flies reared under light, which is suppressed upon the expression of wild type *NECAP* but not by variants associated with diseases. (a. b) Schematic diagrams of Rh1 distribution in photoreceptors. Rh1 mostly localizes to the rhabdomeres of six photoreceptors (R1 to R6) in each ommatidium, organized in a trapezoidal pattern (a). Excessive Rh1 internalization causes additional cytoplasmic Rh1 puncta to appear (b). (c) Immuno-fluorescence labeling of adult *Drosophila* ommatidia with anti-Rh1 (red) and Phalloidin that labels the rhabdomeres (cy5 labeled and pseudocolored in green). The images are from three-day-old flies. Two ommatidia per frame are shown. Note the presence of cytoplasmic Rh1 puncta outside of the rhabdomeres in *NECAP^GFSTF.0^/Y* hemizygous males reared under light (center panels), but not in those reared in dark (right panels). (d) Anti-Rh1 (cy5 labeled in pseudocolored in blue) partially co-localizes with the endosome marker, mCherry-2xFYVE (red). Merged channel (left) and the individual anti-Rh1 (middle) and mCherry channels (right) are shown. Representative areas of co-localization are indicated with white arrows. The genotype is indicated on top of the panels. (e) Immuno-fluorescence images of *NECAP^GFSTF.0^* male ommatidia expressing the indicated transgenes through the *Rh1-Gal4* driver. Anti-Rh1 (red) and Phalloidin (cy5 labeled and pseudocolored in green). Rh1 is mostly within the rhabdomeres when *NECAP^GFSTF.0^/Y* is rescued with the wild type *NECAP* transgene (left), but not with the *NECAP^G176R^* (center) and *NECAP^P19L^* transgenes (right). Scale bars in (c, d, e) represent 1 μm.

We further examined anti-Rh1 signals in the *yw, NECAP^GFSTF.0^/Y* retina that expresses the fluorescent mCherry protein tagged with the endosomal localization sequence, FYVE (McLaughlin et al. 2026) (genotype: *Rh1-Gal4>UAS-mCherry-2xFYVE*). Cytoplasmic anti-Rh1 puncta partially overlapped with the mCherry signal, supporting the endosomal localization of Rh1 in the *NECAP* mutant (Figure 4d).

To confirm that the Rh1 internalization phenotype is due to *NECAP* loss-of-function, we expressed transgenic *NECAP* in the *yw, NECAP^GFSTF.0^/Y* male photoreceptors using the *Rh1-Gal4* driver. Expression of the wild type *UAS-NECAP* transgene suppressed the appearance of Rh1 signals outside the rhabdomeres, but *UAS-NECAP^G176R^* and -*NECAP^P19L^* transgenes failed to suppress the phenotype (Figure 4e). These results further support the idea that *NECAP^GFSTF.0^* is a loss-of-function allele and *NECAP* antagonizes light-dependent Rh1 internalization in photoreceptors.

### *NECAP* mutant adult eyes exhibit abnormal electroretinogram (ERG) patterns

Rh1 at the rhabdomeres responds to light by initiating a phototransduction cascade. The resulting photoreceptor depolarization could be assessed through electroretinogram (ERG) measurements. We compared the Light-Coincident Receptor Potential (LCRP), a component of the ERG that shows the initial depolarization of the photoreceptors in response to a light stimulus, between five-day-old *yw* control flies and *yw, NECAP^GFSTF.0^/Y* males. When exposed to a single stimulus, there was no significant difference in the LCRP amplitude (Figure 5a, b). However, *yw, NECAP* mutant flies exposed to repeated light stimulus show a gradual decline in the LCRP amplitude (Figure 5c, d). This phenotype is pronounced in the null *NECAP^T2A^*, while more subtle in the hypomorphic *NECAP^GFSTF.0^*. This gradual reduction in LCRP could reflect a faster depletion of Rh1 from the apical membrane of the photoreceptors after repeated light stimulus.

**Figure 5:**
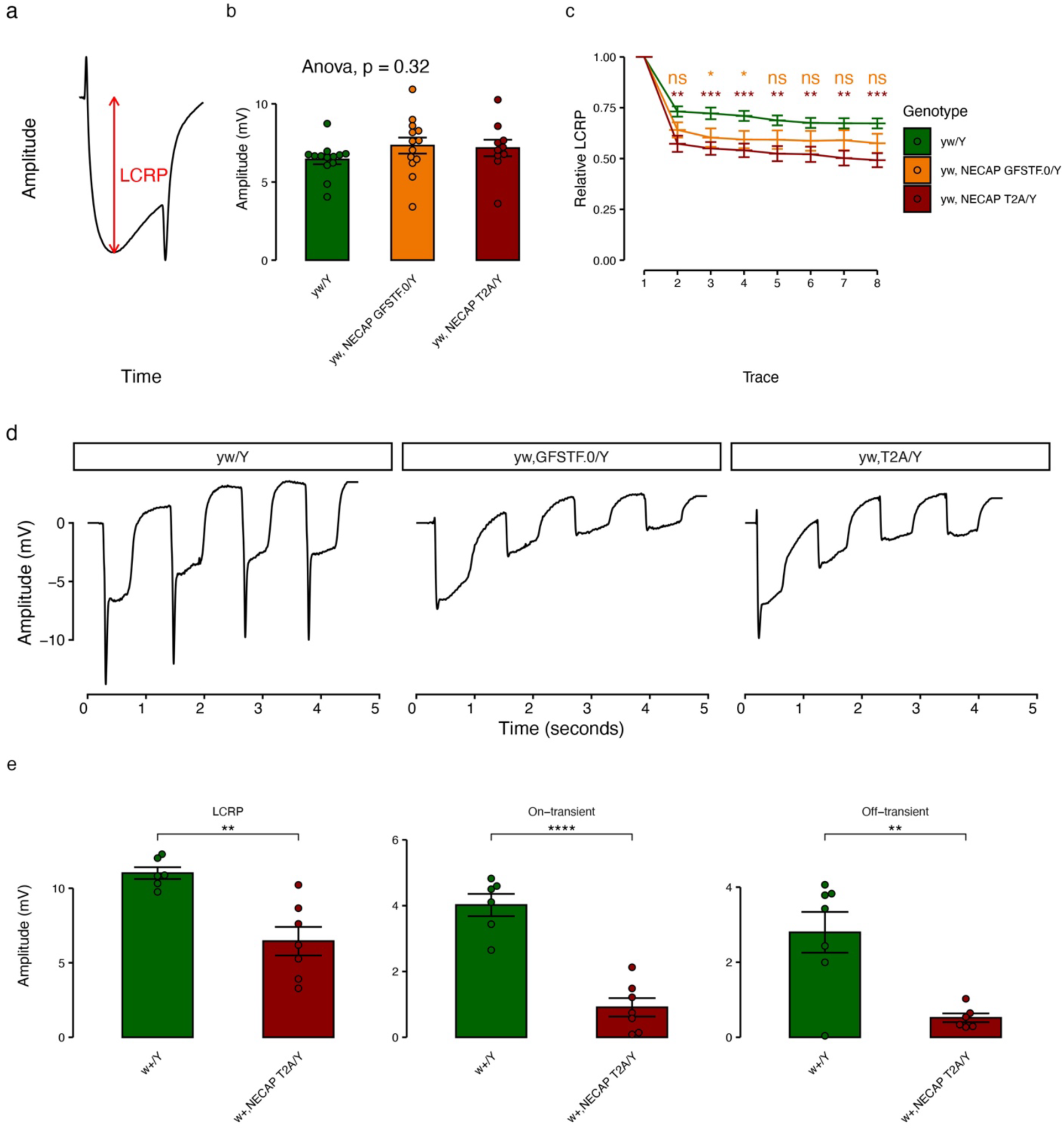
*NECAP* mutant males have an electrophysiological defect in the retina. (a) A typical ERG trace in the fly retina. The LCRP corresponds to photoreceptor neuron depolarization. (b) The LCRP amplitudes in control (genotype: *yw/Y*), and *NECAP* mutant males (genotypes: *yw, NECAP^GFSTF.0^/Y* and *yw, NECAP^T2A^/Y*) are similar when flies are exposed to a single light stimulus. Significance tested by ANOVA. (c) *NECAP^T2A^/Y*, and to a lesser degree *NECAP^GFSTF.0^/Y* flies have reduced depolarization on repeat light stimulation. (d) Example ERG traces in 5 day old flies exposed to repeated light stimuli. (e) LCRP, on-transient, and off-transient measurements from 33 day old flies exposed exposed to individual light stimuli show reduction in all three features of the ERG trace. All flies were reared under constant light. Statistical significance: * p < 0.05, ** p < 0.01,***<0.001,****<0.0001.

Since ERGs of flies in *w^-^* background develop significant variability with age, we recombined the *NECAP^T2A^* allele into a *w^+^* background. We then aged these flies and controls to 33 days of age, where we evaluated individual ERG traces. We found a modest reduction in LCRP and a major reduction in both on- and off-transients in our mutant flies (Figure 5e). This is consistent with a possible neurodegenerative phenotype. Notably, on- and off-transients are due to response of the post-synaptic neuron of the photoreceptor. Therefore, the observed phenotype in our *NECAP* mutant could correspond to a synaptic impairment.

### *NECAP* overexpression suppresses excessive Rh1 internalization and retinal degeneration of *NorpA* mutants

*NorpA* encodes a *Drosophila* homolog of phosphoinositide phospholipase C (PLC) that mediates the visual transduction cascade. The photoreceptors of the null mutant, *NorpA^EE5^*, exhibit massive amounts of Rh1 internalization through CME (Alloway, Howard, and Dolph 2000; Orem, Xia, and Dolph 2006; Kristaponyte et al. 2012). We used this model to test if *NECAP* overexpression is sufficient to antagonize CME of Rh1. As reported by others, the photoreceptors of *NorpA^EE5^* flies reared under light had significant anti-Rh1 signals that do not co-localize with the rhabdomere marker by day five after eclosion, indicative of excessive endocytosis. By 9 days, those ommatidia showed disorganized rhabdomeres, a sign of severe retinal degeneration (Figure 6a). Overexpressing wild type *UAS-NECAP* in this background suppressed these *NorpA^EE5^* phenotypes (Figure 6b). The abnormal distribution of Rh1 in *NorpA^EE5^* mutants was light-dependent, as flies reared in the dark had most Rh1 detected within the rhabdomeres (Figure 6c). *NorpA^EE5^* flies reared with light exposure had reduced total Rh1 levels, consistent with Rh1 undergoing excessive light-dependent endocytosis and ultimate degradation. Overexpression of wild type *UAS-NECAP* suppressed the reduction of total Rh1 in *NorpA^EE5^*mutant flies reared under light (Figure 6d, e). We further assessed the course of retinal degeneration by following Rh1-GFP pseudopupil fluorescence in live adult flies. All *NorpA^EE5^* flies reared under light lost their retinal integrity by 17 days after eclosion (n = 52). But those overexpressing wild type *UAS-NECAP* under otherwise equivalent conditions suppressed the course of retinal degeneration, with 73% of the flies retaining intact pseudopupils at day 17, and 39% with intact pseudopupils at 31 days after eclosion (n = 56). On the other hand, expressing *UAS-NECAP^G176R^*accelerated the degeneration of *NorpA^EE5^* retina, with all flies losing intact pseudopupils by five days after eclosion (n = 31) (Figure 6f), consistent with its dominant negative properties. These results further support the role of *NECAP* in antagonizing light-dependent Rh1 internalization in a well-established model of clathrin-mediated endocytosis.

**Figure 6:**
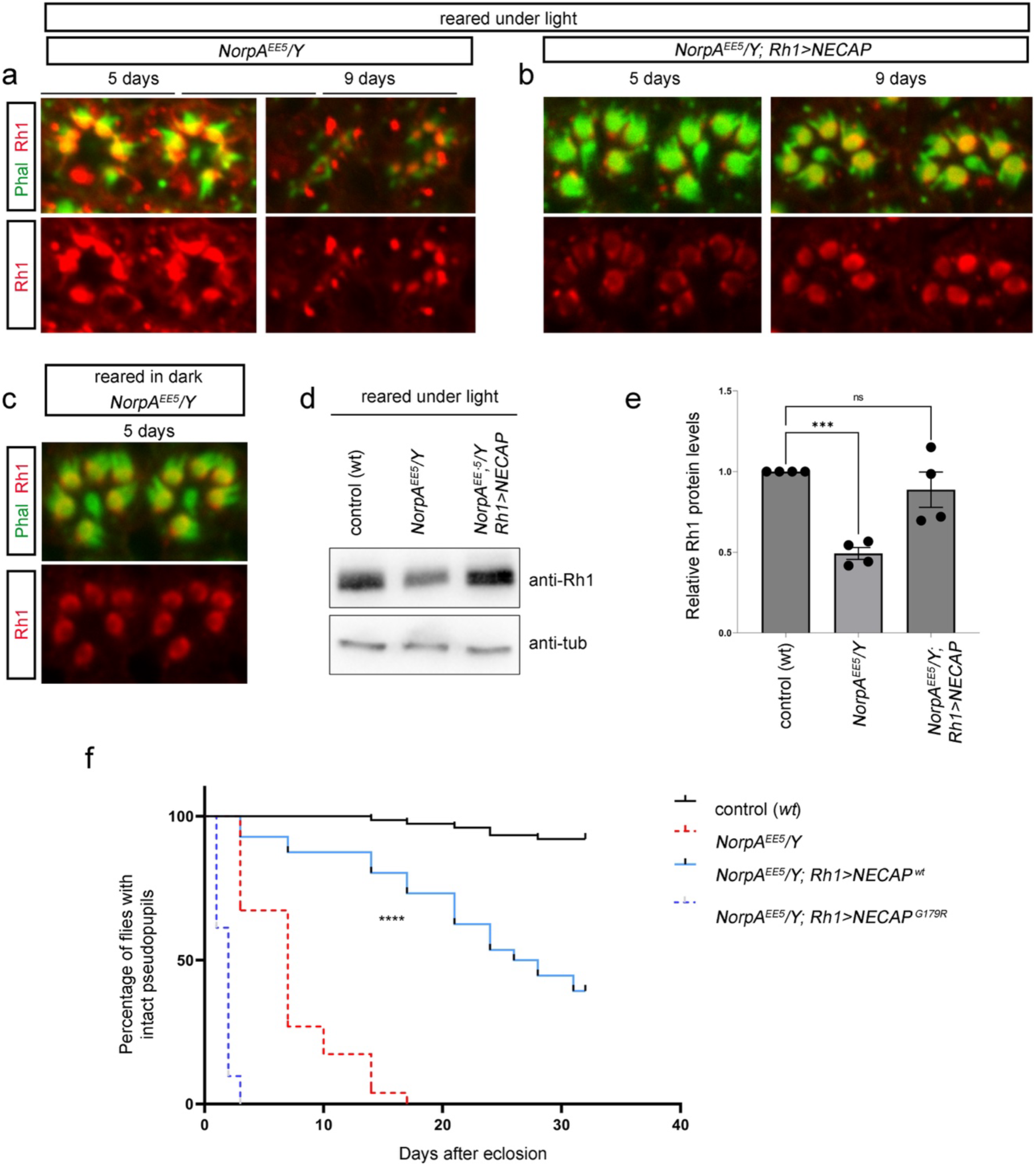
*NECAP* overexpression suppresses excessive Rh1 internalization and retinal degeneration of *NorpA* mutants. (a-c) Immuno-labeling of adult *Drosophila* ommatidia with phalloidin (rhabdomere marker, green) and ant-Rh1 (red). Two ommatidia are shown in each panel. (a) *NorpA^EE5^/Y* males reared under light for the indicated days. (b) *NorpA^EE5^/Y; Rh1-Gal4/UAS-NECAP* flies reared under light for the indicated days. (c) *NorpA^EE5^/Y* flies reared in the dark. (d) Anti-Rh1 western blot of flies of the indicated genotypes reared under light for five days. (e) Quantification of relative anti-Rh1 band intensities, normalized with the anti-tubulin bands. ANOVA followed by Tukey’s post hoc test was used for statistical analysis. N.s. = not significant. *** p < 0.0005. (f) Retinal degeneration as assessed through the percentage of flies retaining intact fluorescent pseudopupils, as assessed with Rh1-GFP. The genotypes are indicated. Male flies were used for the analysis. N numbers for each genotype are as follows: control (*w^1118^*) = 76, *NorpA^EE5^/Y* = 52, *NorpA^EE5^/Y; Rh1-Gal4/UAS-NECAP^WT^* = 56, *NorpA^EE5^/Y; Rh1-Gal4/UAS-NECAP^G176R^* = 31. Log-rank test was used to assess the statistical significance between *NorpA^EE5^/Y* and *NorpA^EE5^/Y; Rh1-Gal4/UAS-NECAP^wt^*. **** p < 0.00005.

## Discussion

NECAP proteins have been characterized as AP-1 and AP-2-interacting proteins, but their precise roles in endosome trafficking and disease have remained unclear. For example, some studies suggest a role of NECAPs in stimulating endocytosis (Ritter et al. 2013; Wrobel et al. 2019), while others support antagonistic roles (Beacham 2018; Partlow 2019). Additionally, mammalian NECAP2 is reported to act independently of CCVs by regulating the AP-1 adaptor complex (Chamberland et al. 2016). Aside from the genetic interactions reported between *C. elegans ncap-1* and *fcho-1* (a muniscin gene) (Beacham 2018), the phenotypic consequences of *NECAP* loss at an organismal level are unclear. Here, we showed that *Drosophila NECAP* is not required for normal development, but the loss-of-function mutants show major phenotypes in photoreceptors. In these cells, *NECAP* specifically antagonizes light-dependent Rh1 internalization, a process that occurs through AP-2-dependent CCVs. We further show that two variants identified in humans and canines are loss-of-function alleles with dominant negative properties.

AP-2 plays diverse roles throughout development, and the loss of AP-2 subunits causes early developmental lethality (Bazinet et al. 1993; González-Gaitán and Stenmark 2003; González-Gaitán and Jäckle 1997; Windler and Bilder 2010). Our *NECAP ^KO^* deletes most of NECAP’s coding sequence, and the *NECAP^T2A^* allele truncates the encoded protein. Still, these mutants still produce viable adults with no obvious morphological defects at adult eclosion. This phenotype suggests that NECAP plays a modulatory role during clathrin-mediated endocytosis, perhaps regulating only a subset of AP-2’s cargo.

We decided to examine the adult retina in part because Rh1 undergoes light-dependent internalization through AP-2-mediated endocytosis. If NECAP is a stimulator of AP-2-mediated endocytosis as reported (Ritter et al. 2013; Wrobel et al. 2019), Rh1 is expected to remain in the rhabdomeres in *NECAP* loss-of-function mutants, whereas an opposite outcome is expected if *NECAP* is an inhibitor of AP-2-mediated endocytosis. The *NECAP* loss-of-function mutants we examined support a role for *NECAP* as an antagonizer of endocytosis. *NECAP* overexpression experiments further substantiate this idea: Expressing wild type *NECAP* transgenes in *NECAP* mutant photoreceptors rescues the Rh1 internalization phenotype. Moreover, *NECAP* overexpression is sufficient to block the massive Rh1 internalization phenotype of *NorpA* mutants, which occurs through the AP-2 and clathrin-dependent endocytosis.

While the phenotype can also be explained by NECAP promoting AP-1-mediated endosome recycling, AP-1-mediated recycling is a function specific for NECAP2 in mammals (Chamberland et al. 2016). Our experiments show that human *NECAP1* and *NECAP2* can both rescue the Rh1 internalization phenotype of *NECAP* mutants, which is more consistent with the model that NECAP antagonizes AP-2-mediated endocytosis.

We were particularly interested in the retinal phenotype because *NECAP1* variants have been associated with canine breeds predisposed to retinal atrophy (Hitti et al. 2019). The cellular nature of the retinal atrophy phenotype in these canine breeds has not been examined. Is it possible that excessive rhodopsin endocytosis, similar to what we found in *Drosophila NECAP* mutants, contributes to retinal atrophy in the canine model? We note that excessive rhodopsin endocytosis caused by certain pathogenic variants of rhodopsin underlie aggressive forms of retinal degeneration in Retinitis Pigmentosa (Chuang et al. 2004; Chen et al. 2006). Moreover, our studies indicate that the canine variant is likely a loss-of-function allele. These observations suggest that excessive rhodopsin internalization is one of the possible mechanisms contributing to retinal atrophy caused by *NECAP1* variants.

In conclusion, we identify *Drosophila NECAP* as a critical regulator of light-dependent Rh1 internalization and photoreceptor survival. While several different mechanisms of NECAP action had been proposed previously, our results are most consistent with NECAP’s in antagonizing AP-2-mediated endocytosis. The mild mutant phenotype suggests some degree of specificity in the CCV cargos regulated by NECAP. Finally, the results suggest a possible pathological mechanism associated with retinal atrophy in mammalian models.

## Materials and Methods

### AlphaFold 3 prediction of NECAP–AP-2mu interaction

Protein complex predictions were performed using the AlphaFold 3 server (Abramson et al. 2024). To model the NECAP–AP-2mu interaction, the PHear domain of mouse NECAP2 (residues 7-139) or *Drosophila* CG9132 (residues 7-135) was analyzed together with the AP-2mu subunit from the corresponding species. For each species, predictions were performed with both the unphosphorylated and phosphorylated forms of AP-2mu (mouse pThr156; *Drosophila* pThr154), with five models generated per condition using default server settings (random seed). All prediction results were similar, and model 0 results are presented here. Predicted complexes were evaluated using iLIS (integrated Local Interaction Score), computed using the AFM-LIS pipeline (Kim 2026) (https://github.com/flyark/AFM-LIS), and ipTM. An iLIS >= 0.223 was used as the threshold for a positive predicted interaction. Structural visualizations were performed using UCSF ChimeraX (Pettersen et al. 2021).

### Fly genetics and husbandry

All fly crosses were maintained at 25 °C, reared with a standard cornmeal-agar diet supplemented with molasses. The following flies used in this study had been reported previously: *Rh1-Gal4* (Mollereau et al. 2000), *Rh1-GFP* (*Rh1* promoter driving GFP) (Pichaud and Desplan 2001), *ninaE-EGFP* (GFP fused to the *Rh1* coding sequence, driven by *Rh1* promoter) (Huang, Xie, and Wang 2015), *NorpA^EE5^* (Yoshioka, Inoue, and Hotta 1985), *UAS-tdTomato* (Fendl, Vieira, and Borst 2020), *UAS-hNECAP1.HA*, *UAS- hNECAP2.HA* (Yamamoto et al. 2014), *UAS-mCherry-2xFYVE* (McLaughlin et al. 2026).

*NECAP^T2A^* and *NECAP^GFSTF.0^* lines were generated in the Bellen lab in the *y, w* genetic background (Li-Kroeger et al. 2018), and are available through the Bloomington Stock Center (stock # 80104, 80105).

The *NECAP^CRISPR^ ^KO^* line was generated at WellGenetics, Inc., with the guide RNAs targeting the following [PAM] sequences:

gRNA1 (upstream): CGCATCCTGCCCGTCCAGGT[GGG]. The cutting site is -12 nt from the ATG of *CG9132-RA/C*.

gRNA2 (downstream): ATAACTCGGCCAACGCCAAC[TGG]. The cutting site is – 15nt from the stop codon of *CG9132*.

These gRNAs caused a 939-bp fragment of *CG9132* to be deleted. The deleted sequence was replaced with a 3xP3-RFP sequence from a knock-in cassette, which has an artificial 3xP3 promoter that drives RFP, flanked by loxP sites. The RFP signal was used to select for successful knock-ins. Genomic PCR was used to validate the insertion at the targeted locus.

To generate *UAS-NECAP* transgenes, wild type and mutant *NECAP* DNA were made through gene synthesis and subcloned into the pUAST-attB plasmid. The transgenic lines were targeted for insertion into the 2^nd^ chromosome by injecting the plasmids into the y, w; PBAC{y[+]-attP-3B}VK00001 ((Venken 2006) Bloomington stock # 9722) using phi31-mediated integration.

For all experiments with flies reared under light, the flies were incubated in the 25 °C incubator with 1000 lux of constant light exposure for day and night. For those experiments with flies reared in dark, the flies were reared in an enclosed cardboard box in the 25 °C incubator.

### Fluorescence Microscopy and Western Blots

Standard protocols were followed for western blots and whole mount immuno-labeling experiments using the following primary antibodies: Mouse monoclonal 4C5 anti-Rh1 (Developmental Studies Hybridoma Bank (DSHB), used at 1:200 for immuno-labeling or retina and 1:5000 for western blots), anti-β-tubulin (Covance #MMS-410P). Rhodamine-conjugated Phalloidin (Molecular Probes cat #R415) was used to image rhabdomeres through whole mount retina labeling.

To image *ninaE-GFP* expressing photoreceptors (Figure 1d, e, S2), live flies were embedded in 2% agarose, and the GFP fluorescence from their retina was imaged directly using Zeiss LSM700 confocal microscope with a 40x water lens.

### Electron Microscopy

Adult flies were anesthetized with CO_2_ and heads were cut in half to facilitate the penetration of the fixative. The samples were incubated in freshly made fixative containing 2.5% glutaraldehyde, 2% paraformaldehyde, and 0.05% Triton X-100 in 0.1M sodium cacodylate buffer (pH 7.2) on a rotator (for 4 hours or more) until all fly eyes sank to the bottom of the tube. The samples were subsequently incubated with an equivalent fixative but without Triton in 4°C for 4 days on rotator. After washing, the fly eyes were post fixed with 1% OsO_4_ for 1.5 hour, dehydrated in a series of ethanol solutions (30%, 50%, 70%, 85%, 95%, 100%), followed by two rinses with propylene oxide before being embedded in EMbed812 epoxy resin (Electron Microscopy Sciences, Hatfield, PA). 500nm thick semi-thin sections were cut, mounted on glass slides and put on a hot plate overnight at 60 °C. The sections were stained with 0.1% Toluidine blue, dried on a hot plate and cover-slipped with Permount mounting medium (Electron Microscopy Sciences, Hatfield, PA) for light microscopy. 70nm ultra- thin sections were cut and mounted on formvar coated slot grids and stained with uranyl acetate and lead citrate. Imaging was performed by an electron microscope (CM12, FEI, Eindhoven, The Netherlands) at 120 kV, and recorded digitally using a camera system (Gatan 4k x 2.7K) with software Digital Micrograph (Gatan Inc., Pleasanton, CA).

Immuno-gold electron microscopy of *Drosophila* ommatidia was done following a previously described protocol (Colley, 1991). Rabbit polyclonal anti-Rh1was used as the primary antibody, and 18nm Colloidal gold anti-Rabbit antibody was used as the secondary.

### Pseudopupil-based retinal degeneration assay

The flies were reared in the 25 °C incubator with 1000 lux of constant light. Retinal degeneration was assessed based on green fluorescent pseudopupils generated by the *Rh1-GFP* transgene. We interpreted clear trapezoidal pseudopupils as evidence in intact photoreceptors, while their disappearance was interpreted as a sign of retinal degeneration.

### PCR

Genomic PCRs to validate the *NECAP^KO^* line (Figure S1) were performed using the following primer pairs:

OWGc1521 (upstream forward primer): CCAACCACAACAATTCGC

OWG0156 (upstream reverse primer): CGAGGGTTCGAAATCGATAA

OWG0167 (downstream forward primer): AACGCAAGCAAATGTGTCAG

OWGc1522 (downstream reverse primer): TTCTGGCCGGTTACACTGAC

RT-PCR results to validate the *NECAP^T2A^* allele (Figure S1) were performed using standard protocols with the following primer pairs:

PCR #1_left primer: TGGCAAGAACAAAGGCTCCT

PCR #1_right primer: TCTGTTGTCCTTGGTTGCCA

PCR #2_left primer: CCAGGAGCAGATCGAGAAGG

PCR #2_right primer: CCCGAGGAGCCTTTGTTCTT

### Electroretinograms (ERGs)

Standard protocols were followed to measure ERG traces (Lauwers and Verstreken 2018). Briefly, a fly was immobilized on a glass slide using glue (Elmer’s). Two glass electrodes were pulled by heating borosilicate glass capillary (Sutter Instrument #BF-120-69-10) to 62℃. These electrodes were then filled with 200 mM saline and affixed to the outside of the retina and the back of the head using micromanipulators. A series of eight 550-lux light pulses was delivered using a halogen light source (AMScope) at a rate of approximately 1/second following 15 seconds for dark acclimation. Electrical activity of the photoreceptor after stimulation was recorded from the electrodes.

### Quantification and statistics

To quantify proteins in gels, we measured average pixel intensities of western blot bands using Image J, and normalized them to anti-β tubulin bands. Graphs were generated after at least three independent measurements and p values were calculated using a one way ANOVA test with a multiple comparisons test. For the retinal degeneration assay in Figure 6, we used the Log-rank (Mantel-Cox) test. Graphs were made using the *Graphpad Prism* program and the R statistical software. All error bars represent SEM (Standard error of the mean).

## Acknowledgements

We thank Gunther Hollopeter for providing comments on the manuscript and Alice Liang at the NYULH DART Microscopy Laboratory (RRID: SCR_017934) for assistance with electron microscopy work. This work was supported by the NIH grant R01 EY020866 to H.D.R. The NYU Langone NYULH DART Core is partially funded by the NYU Cancer Center Support Grant NIH/NCI P30CA016087. N.P. is an investigator of the Howard Hughes Medical Institute. M. T. is supported by the Ting Tsung & Wei Fong Chao Foundation. H.J.B. is supported by the NIH grant R24 OD 031447 and the Huffington Foundation.

## Supplemental Figure Legends

**Figure S1: NECAP sequence alignment and the design of the mutant alleles.** (a) Sequence alignment of NECAP proteins across species. Human NECAP1, NECAP2, *C. elegans* ncap-1, and the two protein isoforms of *Drosophila* NECAP (products of alternative splicing) are aligned. Key amino acids relevant to this study are shown. The *NECAP^KO^* allele used in this study deletes sequences after the 27^th^ amino acid residue in the long protein isoform. The *Drosophila* NECAP protein has high sequence similarity to the human homologs throughout the sequence, including the N-terminal PHear domain and the C-terminal WVQF motif. (b) The design of the *NECAP^KO^* allele. (c) PCR validation of the knockout. The regions amplified by PCR are indicated in (b). (d) The design of the *NECAP^T2A^*allele. (e) PCR validation of the transposon insertion that disrupts transcripts 3’ to the insertion site. The PCR amplified regions are indicated with arrows in (b).

**Figure S2: *NECAP* expression assessed by *yw, NECAP^T2A^*-associated Gal4 driving *UAS-tdTomato* expression.** (a) Whole body adult male flies. The fly on the left is a negative control that has an *FM7* chromosome instead of the *NECAP^T2A^* allele. (b, b’) Adult retina of the genotype *yw, ninaE-GFP, NECAP^T2A^; uas-tdTomato/+* (0 day after eclosion). *ninaE-GFP* expression (green) marks the rhabdomere (apical membrane) of photoreceptor cells. tdTomato fluorescence is detected around those rhabdomeres, where the photoreceptor cell bodies reside. (b’) is the tdTomato only channel of the image shown in (b). Scale bar in b represents 10 μm.

**Figure S3: Immuno-labeling with Phalloidin shows abnormal retinal organization in *NECAP* mutants exposed to constant light.** Confocal microscope images of fixed retina from five-day-old adults reared under constant light. Phalloidin labeling (green) marks rhabdomeres. (a) A control retina (genotype: *yw/w*) shows a regular array of ommatidia, each with seven photoreceptors arranged in a cluster. (b) The *NECAP* homozygous mutant retina (genotype: *yw, NECAP^T2A^/w, NECAP^KO^*) does not have an organized array of ommatidia. The scale bar in b is 10 μm.

